# Bacterial capsular polysaccharides with antibiofilm activity share common biophysical and electrokinetic properties

**DOI:** 10.1101/2022.05.27.493690

**Authors:** Joaquín Bayard-Bernal, Jérôme Thiebaud, Marina Brossaud, Audrey Beaussart, Celine Caillet, Yves Waldvogel, Laetitia Travier, Sylvie Létoffé, Thierry Fontaine, Bachra Rokbi, Philippe Talaga, Christophe Beloin, Noelle Mistretta, Jérôme F.L. Duval, Jean-Marc Ghigo

## Abstract

Bacterial biofilms are surface-attached communities that are difficult to eradicate due to a high tolerance to antimicrobial agents. The use of non-biocidal surface-active compounds to prevent the initial adhesion and aggregation of bacterial pathogens is a promising alternative to antibiotic treatments and several antibiofilm compounds have been identified, including some capsular polysaccharides released by various bacteria. However, the lack of chemical and mechanistic understanding of the activity of these high-molecular-weight polymers limits their use for control of biofilm formation. Here, we screened a collection of 32 purified capsular polysaccharides and identified seven new compounds with non-biocidal activity against biofilms formed by *Escherichia coli* and/or *Staphylococcus aureus*. We analyzed the polysaccharide mobility under applied electric field conditions and showed that active and inactive polysaccharide polymers display distinct electrokinetic properties and that all active macromolecules shared high intrinsic viscosity features. Based on these characteristics, we identified two additional antibiofilm capsular polysaccharides with high density of electrostatic charges and their permeability to fluid flow. Our study therefore provides insights into key biophysical properties discriminating active from inactive polysaccharides. This characterization of a specific electrokinetic signature for polysaccharides displaying antibiofilm activity opens new perspectives to identify or engineer non-biocidal surface-active macromolecules to control biofilm formation in medical and industrial settings.

**Significance statement:** Some bacteria produce non-biocidal capsular polysaccharides that reduce the adhesion of bacterial pathogens to surfaces. Due to a lack of molecular and structural definition, the basis of their antiadhesion activity is unknown, thus hindering their prophylactic use for biofilm control. Here, we identified nine new active compounds and compared their composition, structure and biophysical properties with other inactive capsular polysaccharides. Despite the absence of specific molecular motif, we demonstrate that all active polysaccharides share common electrokinetic properties that distinguish them from inactive polymers. This characterization of the biophysical properties of antibiofilm bacterial polysaccharide provides key insights to engineer non-biocidal and bio-inspired surface-active compounds to control bacterial adhesion in medical and industrial settings.

## Introduction

Bacterial biofilms are widespread surface-attached or aggregated bacteria that can negatively impact human activities when developing on medical or industrial surfaces (1, 2). Due to their high tolerance to antibiotics, biofilms are difficult to eradicate and the prevention of biofilm-associated infections is a major health and economic issue (3, 4). Strategies to prevent biofilm formation often targets the initial steps of bacterial adhesion using surfaces coated by biocidal agents such as broad-spectrum antibiotics or heavy metals (5). These approaches are limited by the rapid masking of the coated surfaces by bacterial and organic debris, and are associated with worrisome selection of resistance upon repeated contact with treated surfaces (6).

Several studies have shown that non-antibiotic anti-adhesion strategies could also efficiently interfere with bacterial biofilm formation (7-14). The design of bio-inspired materials with anti-adhesion surface properties has been proposed to constitute an effective solution to protect patient care equipment, control pathogen colonization, and therefore impede key steps of infection, from initial surface contact to subsequent bacteria-bacteria interactions (15-17). Non-biocidal and bio-sourced strategies are also actively explored, including approaches preventing and/or disrupting biofilms based on the use of quorum sensing inhibitors that interfere with bacterial communications (18). Bacteria also secrete biosurfactants altering material surface properties such as wettability and charge (19, 20). These surface-active compounds reduce surface contacts and contribute to bacterial motility or are involved in competitive interactions between bacteria (12). Whereas many of these molecules correspond to small lipopeptides, recent studies showed that high molecular weight capsular polysaccharides released by various bacteria could prevent adhesion and subsequent cell aggregation and formation of biofilm by a wide range of Gram+ and Gram-bacteria. These include several nosocomial pathogens such as *Escherichia coli, Pseudomonas aeruginosa, Klebsiella pneumoniae, Staphylococcus aureus, Staphylococcus epidermidis* and *Enterococcus faecalis* (7, 9, 11, 21-24).

Unlike secreted bacterial antagonistic macromolecules such as colicins, toxins, phages and a few toxic surface-active compounds (25, 26), these antibiofilm polysaccharides are non-biocidal (11). They impair bacteria-biotic/abiotic surface interactions mediated by adhesion factors such as pili, adhesins or extracellular matrix polymers via the modification of the wettability, charge and overall bacteria-surface contact properties (7, 12, 18). The use of these non-biocidal antibiofilm macromolecules was proposed as a promising resistance-free approach to reduce pathogenic biofilms and biofilm-associated bacterial infections while avoiding side effects caused by broad spectrum biocides (11, 23, 27). However, the limited description of such antibiofilm macromolecules and their associated chemical and structural activities severely hinders their prophylactic use for bacterial biofilm control.

Here, we investigated the antibiofilm activity of a panel of 32 purified Gram+ or Gram-bacterial capsular polysaccharides of known composition and structure. Among those, we identified nine new non-biocidal polysaccharides inhibiting biofilm formation by prototypical nosocomial pathogens, including *E. coli* and *S. aureus*. This enabled us to perform a structure-function comparison of enough active and inactive polysaccharides and to show that antibiofilm capsular polysaccharides are characterized by a high intrinsic viscosity and a specific electrokinetic signature. Our study therefore identified key chemical and biophysical properties of bacterial antibiofilm polysaccharides, providing insights paving the way for engineering of new and well-defined non-biocidal surface-active macromolecules to control biofilm formation in medical and industrial settings.

## Results

### Size integrity is a key parameter of antibiofilm polysaccharide activity

To investigate the potential relationships between composition, structure, size and activity of non-biocidal bacterial antibiofilm polysaccharides, we first determined the structure of Group 2 capsule (G2cps), a previously identified hydrophilic and negatively charged antiadhesion polysaccharide active on both Gram+ and Gram-bacteria, which is naturally produced and released, notably, by uropathogenic *E. coli* strains (7). G2cps was purified by High-Performance Anion-Exchange Chromatography – Pulsed Amperometric Detection (HPAEC-PAD), High Performance Size-Exclusion Chromatography coupled to Static Light Scattering (HPSEC-LS) and analyzed by ^1^H and ^13^C nuclear two dimensional ^1^H- ^31^P experiment magnetic resonance (NMR) studies. These analyses showed that G2cps polysaccharide consists of repeating units composed of O-acetylated and glycerol phosphate residues with an average molecular weight of 800 kDa as illustrated in Figure 1A. This structure is similar to that of *E. coli* polysaccharides K2 and K62 (also named K2ab) (28). We then gradually reduced the size of G2cps polysaccharide while preserving its structural integrity using radical oxidation hydrolysis (Supporting Figure S1). The determination of antibiofilm activity of full length and fragmented G2cps showed that even minor reduction of polysaccharide size resulted in a loss of G2cps activity (Figure 1B), indicating that the conservation of the size of the G2cps polymer is critical for its antiadhesion properties.

**Figure 1.**
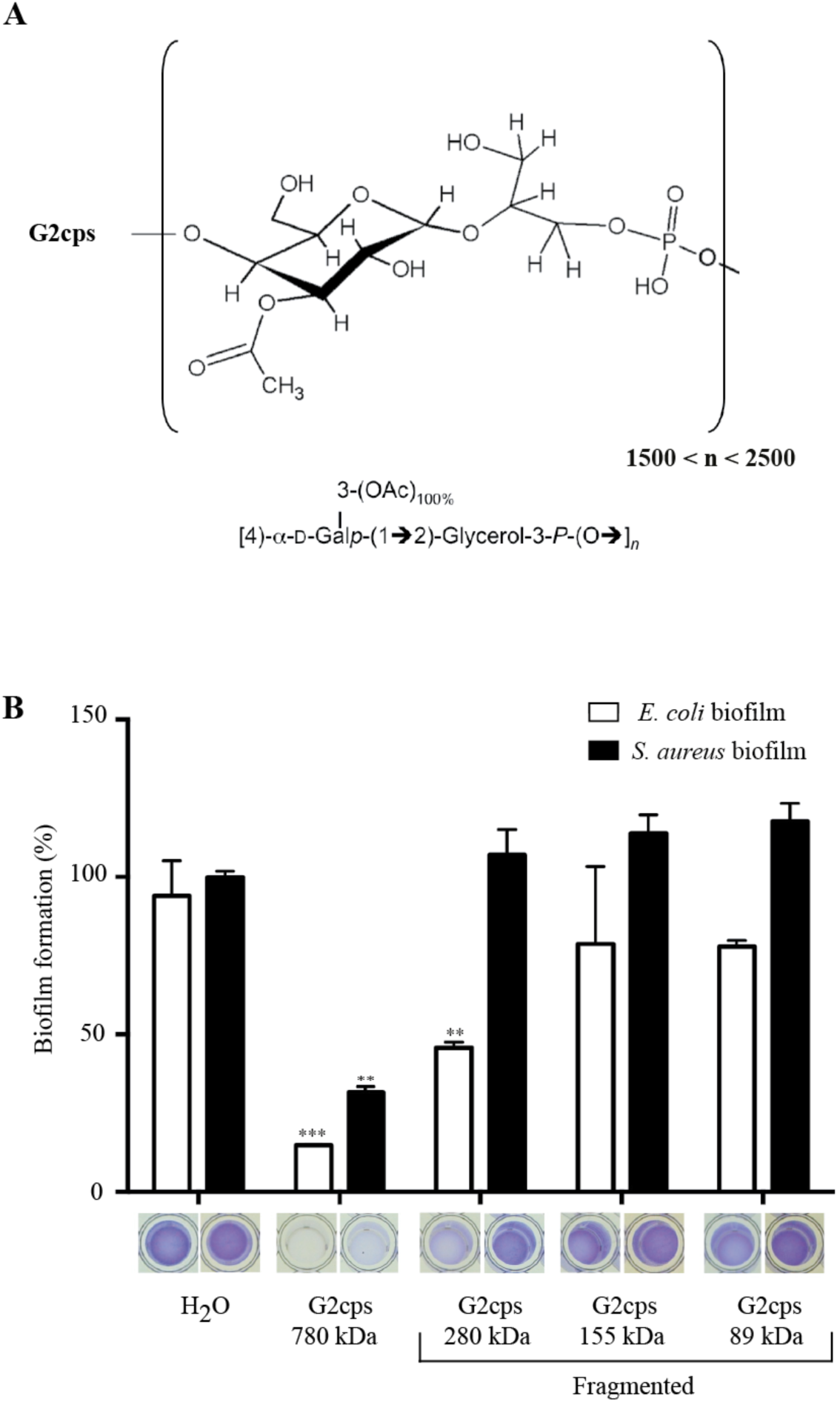
Structure of the G2cps polysaccharide and its antibiofilm activity as a function of the extent of its fragmentation. **A**. Structure and composition of G2cps polysaccharide. **B**. Biofilm inhibition activity against *E. coli* and *S. aureus* biofilm of native and fragmented G2cps polysaccharide. Biofilm assays were performed in the presence of 50 μg/ml of polysaccharide. Each experiment was performed at least 3 times. ** p<0.01; *** p<0.001.

### Screening a collection of bacterial capsular polysaccharides reveals new non-biocidal antibiofilm compounds

To further attempt to identify G2cps structural or composition features associated with its activity, we screened a collection of 30 different high molecular-weight bacterial polysaccharides to detect new macromolecules with G2cps-like antibiofilm properties. Most of them are capsular polysaccharides of known origin, composition and structure produced and purified from different strains of *Streptococcus pneumoniae, Salmonella enterica* serovar Typhi, *Haemophilus influenzae, Neisseria meningitidis*, and used as antigens in several polysaccharide and glycoconjugate human vaccines (Supporting Table S1). We showed that, at equal concentration (100µg/mL), Vi, MenA and MenC polysaccharides were as active as G2cps in inhibiting *E. coli* biofilm formation but were less active on *S. aureus* biofilms (Figure 2A, B). Moreover, whereas PRP and PnPS3 were active on both bacteria, the activity of PnPS12F was restricted to *E. coli* and that of PnPS18C to *S. aureus* (Figure 2A, B). The comparison of the activity of Vi, PRP, PnPS3, MenA and MenC showed that Vi is the most active polysaccharide and all 5 polysaccharides are non-biocidal (Supporting Figures S2 and S3). Finally, we confirmed the activity of Vi, MenA, PRP and PnPS3 on a panel of biofilm-forming Gram+ or Gram-bacteria, including a biofilm-forming *E. coli* K-12 MG1655 carrying a F conjugative plasmid, *Enterobacter cloacae* 1092, *K. pneumoniae* Kp21, *S. aureus* 15981 and *S. epidermidis* O47 (Supporting Figure S4).

**Figure 2.**
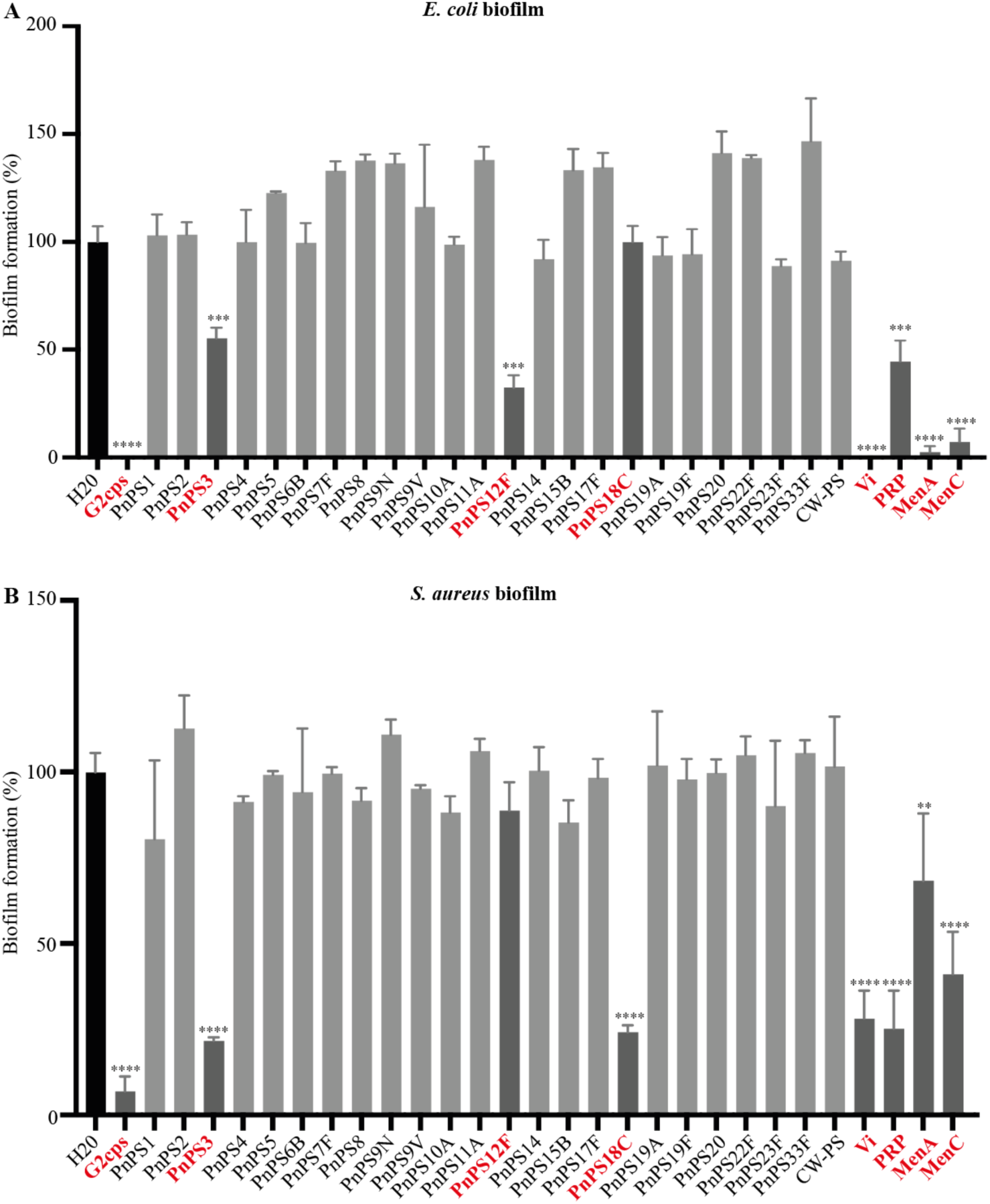
Antibiofilm activity of a collection of bacterial polysaccharides. *E. coli* (panel **A**) *or S. aureus* (panel **B**) biofilm formation assay in the presence of purified bacterial polysaccharides. G2cps was included as a positive control of active macromolecules and water was used as negative control. Red : active polysaccharides with Vi, MenA, MenC, G2cps, PRP and PnPS3 displaying broad spectrum of action, while PnPS18C is only active against Gram+ bacteria and PnPS12F against Gram-bacteria. Experiments were performed twice and in triplicate in presence of 100 μg/mL of each macromolecule. ** p<0.01; *** p<0.001; **** p<0.0001.

### Polysaccharide conformation and high intrinsic viscosity are predictive indicators of antibiofilm activity

To identify the structural and biophysical properties that potentially correlate with polysaccharide antibiofilm activity, we determined the molecular weight (*Mw*) and the intrinsic viscosity ([*η*]) of active (Vi, MenA, MenC, PnPS3, PRP, G2cps) and inactive (PnPS1, PnPS8, PnPS19A, PnPS9V, PnPS7F, PnPS22F and PnPS14) polysaccharides by HPSEC (Table 1). Intrinsic viscosity [*η*] of a given polysaccharide reflects its contribution to the viscosity *η* of the whole solution, i.e. [*η*]=(*η−η*_*0*_)/(*η*_*0*_ *ϕ*), where *η*_*0*_ is the solution viscosity in the absence of polysaccharide and *ϕ* stands for the volume fraction of polysaccharides in solution. Accordingly, [*η*] depends on the conformation adopted by the polysaccharides in solution. This conformation being itself mediated by several physicochemical parameters, including the electrostatic charges carried by the polysaccharide and the charge distribution within the macromolecular body. The combination of *Mw* and [*η*] parameters are indicative of the volume (per mass unit) occupied by the polysaccharides in solution. This analysis revealed a remarkable correlation between intrinsic viscosity and broad-spectrum antibiofilm activity. Indeed, all inactive macromolecules systematically display the lowest, whereas active polysaccharides were characterized by a high (> 7dl/g) intrinsic viscosity (Supporting Figure S5). Moreover, molecular weight *Mw* and intrinsic viscosity [*η*] of polysaccharides with intermediate narrow-spectrum activity (PnPS18C and PnPS12F) cover range of values measured for both broad-spectrum active and non-active macromolecules. To determine whether high intrinsic viscosity could be indicative of a potential antibiofilm activity, we screened additional purified bacterial capsular polysaccharides of our collection and we identified two such non-biocidal polysaccharides, MenY and MenW135, presenting high intrinsic viscosity (Table 1 and Supporting Figures S3 and S5). Although MenY and MenW135 differ in their primary composition from Vi, MenA, MenC and G2cps, they both exhibited similar broad-spectrum antibiofilm activity (Figure 3 and Supporting Figure S6). These results indicated that specific polysaccharide conformation, reflected by a high intrinsic viscosity (29), is a determinant of antibiofilm activity.

**Table 1:**
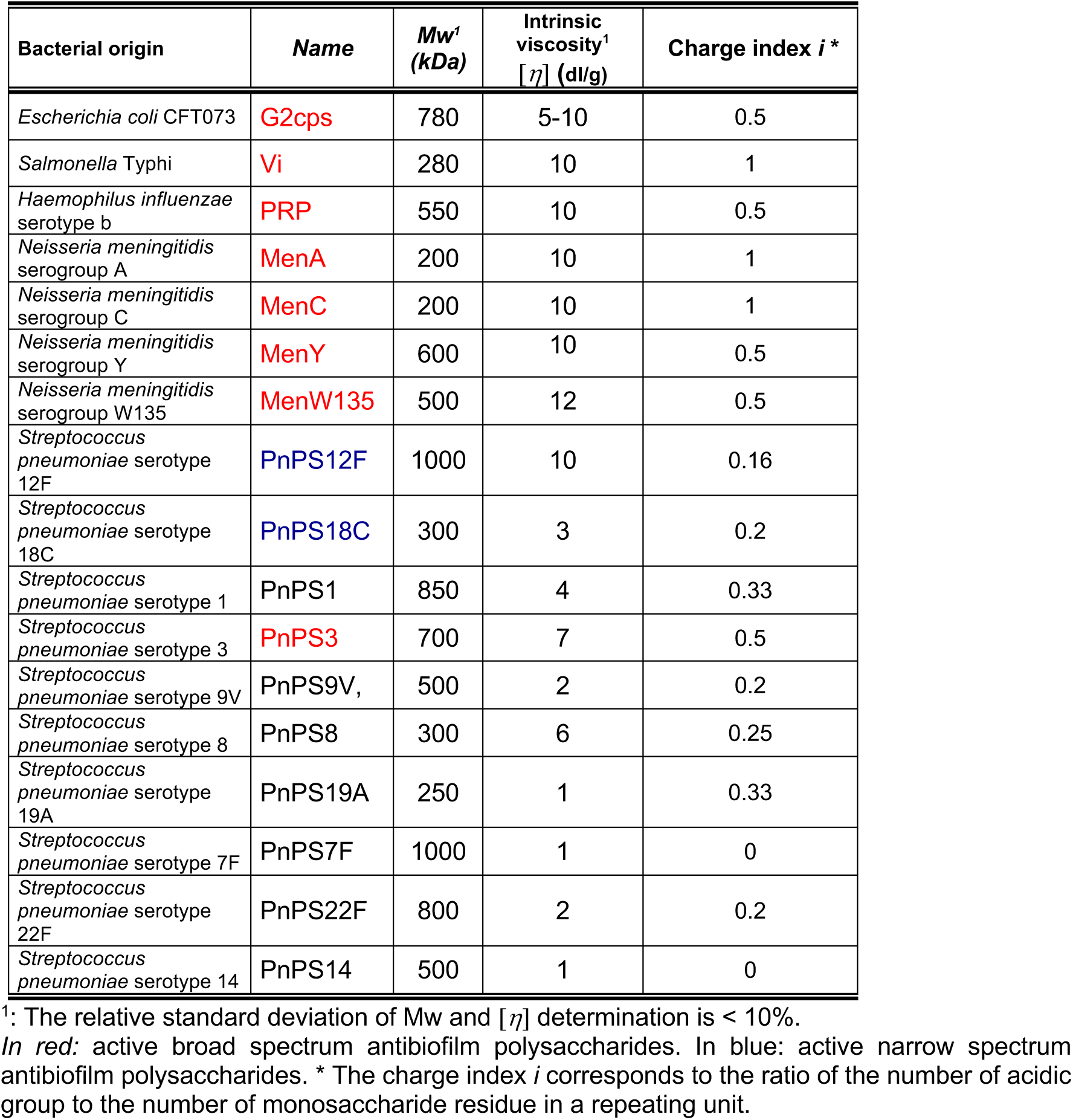
Relevant origin, and physicochemical properties of the polysaccharides used in this work. Indicated values correspond to mean values.

**Figure 3.**
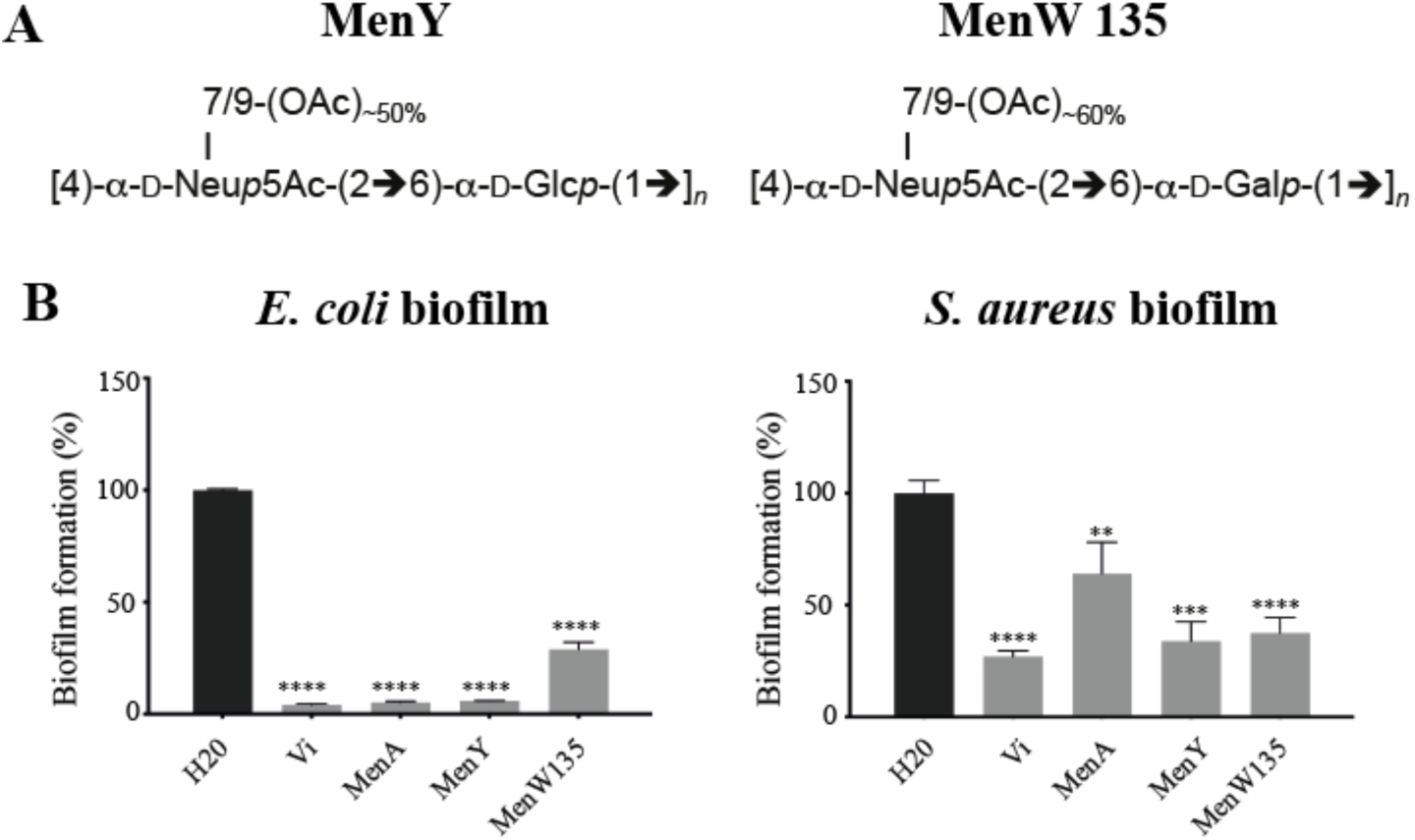
MenA and MenY antibiofilm activity. **A**. Chemical composition and acidic/hydrophobic group ratio of the *N. meningitidis* polysaccharides MenY and MenW135. **B**. Antibiofilm activity of MenY and MenW135 in comparison with previously identified active macromolecules. Biofilm inhibition tests were performed in presence of 50 μg/mL of polysaccharide. Distilled water was used as a negative control. Each experiment was performed at least 3 times. ** p<0.01; *** p<0.001; **** p<0.0001.

### Active antibiofilm polysaccharides are highly negatively charged macromolecules

To further identify the molecular bases of the biophysical properties displayed by active polysaccharides, we first compared the composition of the broad-spectrum polysaccharides (Vi, MenA, MenW, MenY, MenC, G2cps, PRP, PnPS3) but could not identify a chemical feature common to all active molecules. Alternatively, we tested whether an increased surface hydrophilicity, previously associated with G2cps polysaccharide properties (7, 9), could correlate with the antibiofilm activity of the newly identified active polysaccharides. However, the determination of the surface contact angle of a drop of water on hydrophilic glass and hydrophobic plastic surfaces treated with either distilled deionized water or active and inactive polysaccharides did not reveal any correlation between surface hydrophilicity/hydrophobicity balance and antibiofilm activity (Supporting Figure S7). As an additional support of this finding, the characterization of polysaccharide-coated surfaces by Atomic Force Microscopy (AFM) operated in force spectroscopy mode using CH_3_-functionalized nanometric tips did not reveal any correlation between antibiofilm activity, structural surface patterns of adsorbed polysaccharides and surface hydrophobicity/hydrophilicity (Supporting Figure S8). We also evaluated the macromolecular charge, defined by a charge index *i* corresponding to the ratio between the number of acidic groups (carried by uronic and neuraminic acid groups or by phosphate groups) and the number of monosaccharide residues in a repeating unit (Table 1). The determination of this charge index showed that all active polysaccharides (Vi, MenA, MenW135, MenY, MenC, G2cps, PRP, PnPS3) correspond to highest *i*-values with *i* = 0.5 or 1. By contrast, macromolecules without anti-biofilm activity and with narrow-spectrum activity could not be differentiated based on their respective *i*-values. For instance, PnPS18C (narrow activity) for instance, shares similar value of *i* (= 0.2) with PnPS9V and PnPS22F (no activity). These results suggested that the density of negative charges carried by the polysaccharides could be a key determinant of the biophysical properties associated with broad-spectrum antibiofilm activity.

### Active and inactive antibiofilm polysaccharides display distinct electrokinetic patterns

Previous reports documented the link between structure, surface charge organization and electrokinetic properties of macromolecules (30-32) as derived from electrophoresis. Bacterial polysaccharides are paradigms of ‘soft’ macromolecules, i.e. polyelectrolytic assemblies defined by a 3-dimensional charge distribution, permeable to ions from the background electrolyte solution and to the electroosmotic flow developed under electrophoresis measuring conditions (30-32).

To determine whether the electrokinetic properties of polysaccharides (which includes both charge density and flow permeability features) are linked or not to their anti-biofilm activity, we performed blind measurements of the electrophoretic mobility (*μ*) as a function of NaNO_3_ electrolyte concentration in solution (1mM to 100mM range) for eight active, broad spectrum (Vi, MenA, MenW135, G2cps, PRP, PnPS3) or narrow spectrum (PnPS18C and PnPS12F) polysaccharides. The electrophoretic mobility of these active polysaccharides was then compared to those of seven inactive polysaccharides (PnPS1, PnPS9V, PnPS8, PnPS19A, PnPS7F, PnPS22F and PnPS14) (Figure 4). All tested polysaccharides displayed the characteristic electrophoretic signature expected for soft macromolecules, with an electrophoretic mobility tending to a constant, non-zero mobility plateau value (noted below as *μ*\*) at sufficiently large salt concentrations (>30 mM). This property is the direct consequence of the penetration of the electroosmotic flow within the charged polysaccharide globular structure (33). Further qualitative inspection of the sets of electrokinetic data collected for the polysaccharides of interest revealed two main electrokinetic patterns. The first one corresponded to the 7 tested inactive macromolecules whose electrophoretic mobility systematically tends to value of *μ*\* satisfying 0.5< │*μ*\*│ <1.5×10^−8^ m^2^ V^−1^ s^−1^. For these macromolecules, the absolute value of the electrophoretic mobility decreases with increasing electrolyte concentration as a result of screening of the polysaccharide charges by the electrolyte ions. This feature is also shared by the active macromolecules (PnPS3, PRP and G2cps) with the noticeable difference that their asymptotic mobility value │*μ*\*│ is significantly larger with *μ*\* now satisfying the inequality │*μ*\*│ >2×10^−8^ m^2^V^−1^s^−1^ (Figure 4A). The second observed electrokinetic pattern applies to macromolecules with narrow (PnPS18C and PnPS12F) or broad-spectrum (Vi, MenA, MenW135) activities and for which the electrophoretic mobility *μ* poorly depends on background electrolyte concentration. Strikingly, active macromolecules with narrow spectrum antibiofilm activity are defined by electrophoretic mobilities (│*μ*\*│ < 0.5×10^−8^ m^2^ V^−1^ s^−1^) that are much lower in magnitude compared to those measured for broad-spectrum antibiofilm polysaccharides (MenA, MenW135 and Vi, │*μ*\*│ > 2 ×10^−8^ m^2^ V^−1^ s^−1^) (Figure 4B). These results demonstrated clearly that active capsular polysaccharides are characterized by a specific electrokinetic signature.

**Figure 4.**
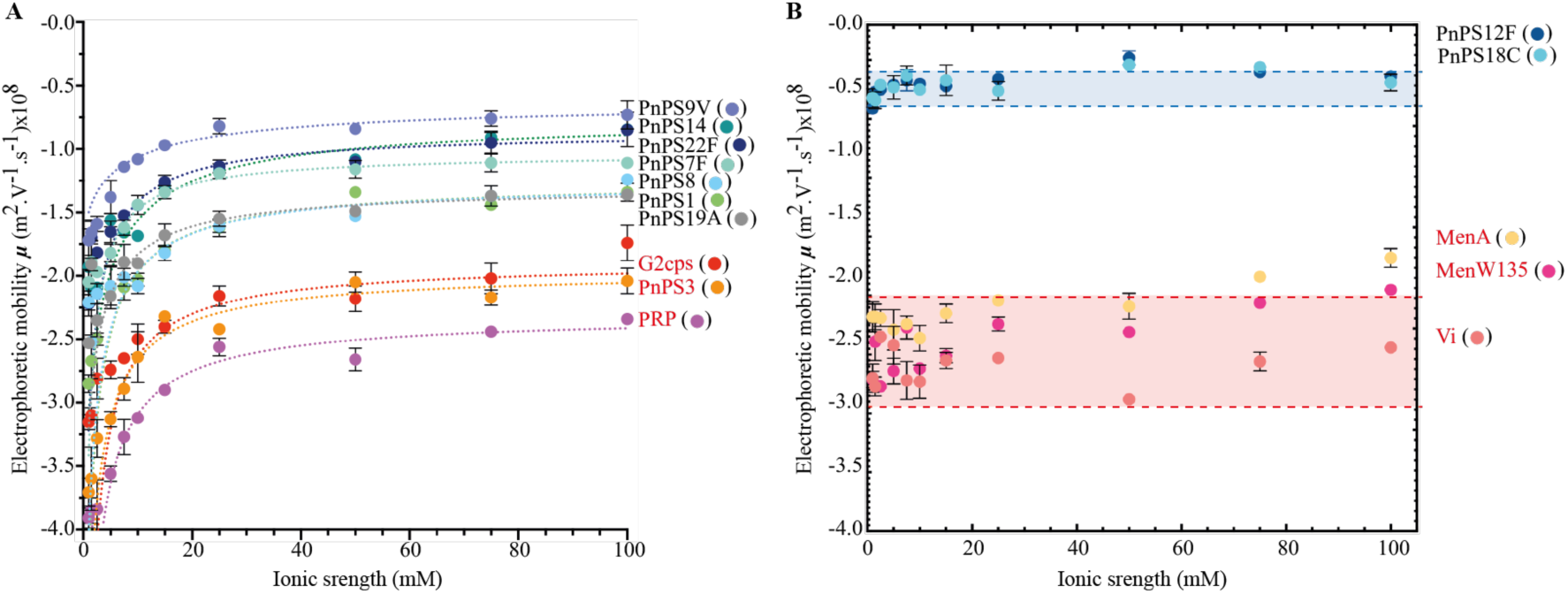
Dependence of the electrophoretic mobility of polysaccharides on NaNO_3_ electrolyte concentration. Points: experimental data. Dashed lines: theory (eq 1). **A**: Polysaccharides whose electrophoretic mobility significantly depends on solution ionic strength (fixed by NaNO_3_ electrolyte concentration). **B**: Bacterial polysaccharides whose electrokinetic features poorly depend on electrolyte concentration. Polysaccharide with name in red display a broad-spectrum antibiofilm activity. PnPS12F and PnPS18C display a narrow spectrum antibiofilm activity (see Table S1). In panel B, the blue zone delimited by blue dotted lines bracket the quasi-electrolyte concentration-independent electrophoretic mobility values measured for the polysaccharides with narrow-spectrum activity. The red zone delimited by red dotted lines bracket the quasi-electrolyte concentration-independent electrophoretic mobility values measured for the polysaccharides with broad spectrum activity.

### The electrokinetic signature of antibiofilm polysaccharides is associated to their high flow permeability and high density of electrostatic charges

To further explore the electrokinetic properties of active capsular polysaccharides we interpreted quantitatively the dependence of the electrophoretic mobility (*μ*) of the tested macromolecules on background electrolyte concentration with soft surface electrokinetic theory. This theory was developed by Ohshima for the electrophoresis of soft colloids (33) in the Hermans-Fujita’s limit that is applicable to soft polyelectrolytes (34) (eq 1 in Materials and Method section). To that end, the required hydrodynamic radii of the macromolecules of interest were determined by Dynamic Light Scattering (DLS) after conversion of the measured diffusion coefficients by means of Stokes-Einstein equation (See Methods and Supporting Table S2). This quantitative analysis highlighted an excellent reconstruction of the electrophoresis data measured for all tested macromolecules tested (Figure 4) and, most importantly, a remarkable classification of their activity (Figure 5) according to their electrostatic and flow permeability properties.

**Figure 5.**
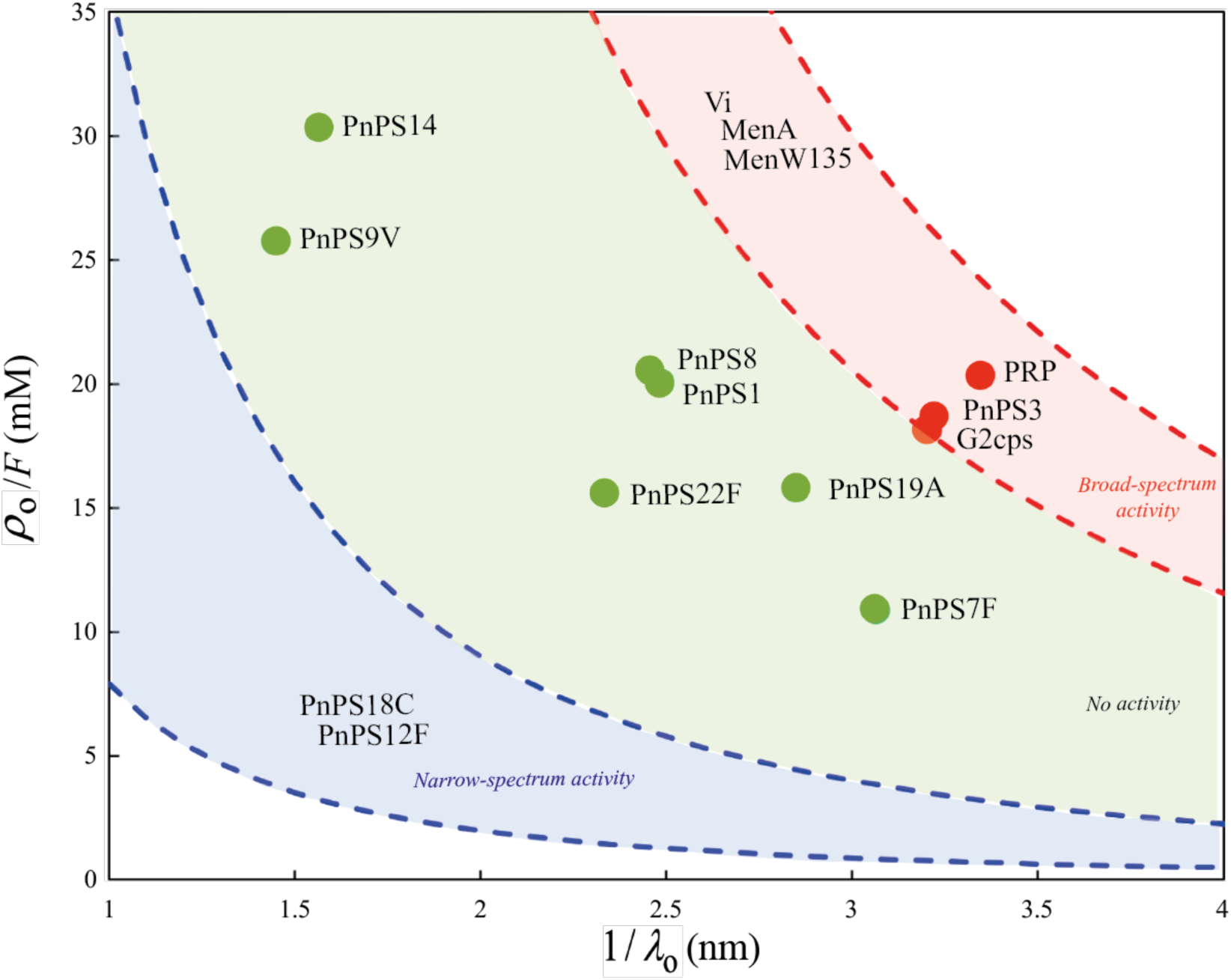
Classification of the antibiofilm activity of the tested polysaccharides according to their charge density *ρ*_o_ and flow penetration length scale 1/*λ*_o_. The charge density *ρ*_o_ (in C m^−3^) is given here in the form of an equivalent concentration of anionic charges defined by *ρ*_o_ /*F* (in mM) with *F* the Faraday constant. The couple (*ρ*_o_,1/*λ*_o_) associated to each macromolecule is retrieved from the modelling of the dependence of measured electrophoretic mobility on the electrolyte concentration in solution according to eq 1. The dotted lines correspond to the set of solutions (*ρ*_o_,1/*λ*_o_) to the equation *µ=µ*\*obtained for the macromolecules PnPS18C and PnPS12F and for the macromolecules Vi, MenW135 and MenA (see colored parts in Figure 4B) whose mobility *µ* over the whole range of electrolyte concentrations does not significantly deviate from the mobility value *µ*\* measured at large electrolyte concentrations (100 mM). For these macromolecules, the poor dependence of *μ* on electrolyte concentration renders difficult any accurate evaluation of both *ρ*_o_, and 1/*λ*_o_ as electrokinetic data fitting then reduces to solve one equation (eq 1) with two unknowns (*ρ*_o_ and 1/*λ*_o_). The space of solutions (*ρ*_o_, 1/*λ*_o_) associated with Vi, Men135, MenA and PnPS18C, PnPS12F correspond to the red and blue zones, respectively. They are delimited by dotted lines that correspond to the (*ρ*_o_, 1/*λ*_o_) solutions of the equation *µ=µ*\* where *µ*\* identifies with the polysaccharide mobilities marked by the dotted lines represented in Figure 4B.

These properties are expressed by two key parameters retrieved from theoretical fitting of electrophoretic data to eq 1, namely: *ρ*_*o*_, which is the net density of negative charges carried by the macromolecule, and *λ*_*o*_ the so-called softness parameter. 1/*λ*_*o*_ (also called the Brinkman length) corresponds to the extent of penetration of the electroosmotic flow within the macromolecule. 1/*λ*_*o*_ is intimately correlated to the polysaccharide structural compacity and by the degree of entanglement of its constituting chains. All macromolecules with narrow spectrum activity (PnPS18C and PnPS12F) are in the lower left zone within the diagram reporting the determined value of *ρ*_*o*_ as a function of the corresponding 1/*λ*_*o*_ (Figure 5). By contrast, polysaccharides with a broad-spectrum activity (e.g. Vi, MenA, MenW135, PRP, PnPS3, G2cps) combine high charge density and high Brinkman length scale, and they are thus found at the upper right region in the *ρ*_*o*_*-*1/*λ*_*o*_ representation (Figure 5). All tested inactive macromolecules (PnPS1, PnPS9V, PnPS8, PnPS19A, PnPS7F, PnPS22F and PnPS14) are positioned in a region intermediate between those of the broad-spectrum active and narrow-spectrum active macromolecules. Taken together, our results indicate that despite the lack of specific molecular motif associated with antibiofilm properties, the combination of a loose structure (i.e. a large permeability to flow due to a large amount of intraparticulate voids) and a high density of carried electrostatic charges is critical for polysaccharides to exhibit antibiofilm activity.

## DISCUSSION

The inhibition of bacterial adhesion using surface-active compounds is considered a promising approach to prevent the key initial steps of bacterial biofilm formation. In this study, we identified nine new non-biocidal antibiofilm macromolecules among a collection of 30 different high molecular-weight bacterial capsular polysaccharides of known structure and composition. This allowed us to compare their chemical, structural and electrokinetic properties to identify key molecular and biophysical determinants discriminating active polysaccharides from inactive ones.

The identified active polysaccharides are composed of a high number of repeating oligosaccharides with different composition and structure. This suggests that, beyond the specific composition of the active polysaccharide, the determinant of their antibiofilm activity could lie in higher order molecular features. Consistently, we showed that, in the case of G2cps, the size integrity of the polysaccharide is critical to maintain antibiofilm activity. We also demonstrated that a high intrinsic viscosity is a property shared by broad spectrum active macromolecules, which displayed the highest charge index *i* among all tested polysaccharides tested. The hydrodynamic radii measured for all tested macromolecules was shown to cover a relatively narrow range, between ca. 15 and 40 nm in hydrodynamic diameter (Supporting Table S2). Such small variations in particle size cannot account for the well differentiated range of intrinsic viscosity measured for non-active and active macromolecules (1 to 6 and 7 to 12 dl/g, respectively, see Supporting Figure S5), especially since both active and non-active polysaccharides display identical molecular weight ranges (Supporting Figure S5). In contrast, the high intrinsic viscosities measured for active macromolecules correlate with their charge index *i*, indicative of a high charge density. This correlation could be due to the fact that viscosimetric properties of particle dispersions depend on their electrostatic characteristics that govern the extent of so-called primary and secondary electroviscous effects (35, 36). These effects are a direct consequence of the presence of charged electric double layers (EDL) at the macromolecule/solution interface and of ensuing particle-particle and particle-fluid electrohydrodynamic interactions: the larger the density of particle charges, the more significant electro-viscous effects become (38,39). As a consequence, the viscosity of a solution containing macromolecules of similar size can significantly differ according to the density of their carried electrostatic charges (35, 36). A large flow penetration within the particles body (as revealed by a high Brinkman length, 1/*λ*_*o*_, see Figure 5) is consistent with the existence of a relatively loose structure adopted by the macromolecules and a reduced frictional force they exert on electro-osmotic flow during electrophoresis, with a resulting significant electrophoretic mobility. Altogether, the identified properties of loose macromolecular structure combined with a high charge density and a related high intrinsic viscosity correlate with antibiofilm activity.

Analysis of the electrokinetic data revealed very distinct electrokinetic patterns associated with broad and narrow spectrum of antibiofilm activities, whereas inactive macromolecules exhibited an intermediate electrokinetic behavior (Figure 5). This suggests that the high and low magnitudes of the polysaccharide charges could determine their broad and narrow activity, respectively, since a charge density with magnitude lying in between these two extremes led to a loss of activity. However, we previously showed that, despite its high negative charge, G2cps displayed low affinity for cationic dyes (7), suggesting that its interaction with surrounding biotic or abiotic environment is not only driven by electrostatics, but may also include remodeling of its surface properties. This could possibly include changes in surface hydration or steric repulsion, and subsequent limitation of bacterial adhesion (7). This indicates that consideration of polysaccharide electrostatic properties alone is not enough to account for their antibiofilm activity. Consistently, analysis of our electrokinetic data sets suggests indeed that the very organization of the polymer chains and the resulting flow permeability properties of the polysaccharidic macromolecules (as qualitatively indicated by the magnitude of their Brinkman length) play an important role in defining their antibiofilm activity. Considering that only charges located in the peripheral region of the macromolecules are probed by electrophoresis (35), the location of these charges and their degree of exposition to the outer solution are likely important factors underlying antibiofilm activity.

Then, how does polysaccharide molecular composition correlate with antibiofilm activity? Most active macromolecules display a high density of negative charges that could contribute to the electrostatic repulsion of negatively charged bacteria. However, bacterial surface structures such as pili, fimbriae, lipopolysaccharides or even poly-cationic exopolysaccharides are known to overcome these repulsive forces and promote bacterial adhesion and aggregation (37-39). We therefore hypothesize that the directionality of the interactions on surfaces (40) could be a critical determinant of broad spectrum antibiofilm polysaccharide activity. The proper exposition of electrostatic charges could optimize the steric repulsion effect in the antibiofilm macromolecules identified in this study. In addition, their high intrinsic viscosity and relative loose structure could mechanically modify the local conditions of adhesion and alter the perception of the surface by bacteria, thereby minimizing their adhesion in the presence of the identified polysaccharides (41). We therefore propose that the active antibiofilm polysaccharides identified in our study could have multi-pronged activity, with both short and long-range modifications of the surface-bacteria or bacteria-bacteria interactions, in particular via an alteration of the local adhesion conditions prevailing on the adhesion surface. The distinction between broad, narrow or lack of activity would therefore depend on the specific combination of biophysico-chemical properties displayed by each macromolecule.

By providing a better definition of the chemical and structural basis of the broad-spectrum antibiofilm activity displayed by capsular polysaccharides, our study offers the possibility to identify new surface-active polysaccharides on the basis of their biophysical properties but also to design and engineer macromolecules mimicking antibiofilm polysaccharide activity through total or partial synthesis. These molecules could be used as non-biocidal biofilm control strategies in prophylactic treatment against the initial adhesion of biofilm-forming pathogens developing on medical and industrial materials.

## MATERIALS AND METHODS

### Bacterial strains and growth conditions

Bacterial strains used in this study are listed in Supporting Table S3. Gram-bacteria were grown in 0.4% glucose-M63B1 minimal medium (M63B1) or in lysogeny broth (LB) medium at 37°C, with appropriate antibiotics when required. Gram+ strains were cultured in tryptic soy broth (TSB) supplemented with 0.25% or 0.5% glucose (*E. cloacae*). Potential biocidal effect of antibiofilm polysaccharides was evaluated from growth curve determination in the presence of 100 μg/ml of purified polysaccharide.

### Bacterial polysaccharides

*S. pneumoniae, N. meningiditis, S*. Typhi *and H. influenzae* capsular polysaccharides used in this study were obtained from Sanofi, Marcy l’Etoile France and Swiftwater, USA. *E. coli* polysaccharides were produced at the Institut Pasteur, Paris. Teichoic acid polysaccharide (cell wall PS, CWPS) from *Streptococcus pneumoniae* strain CSR SCS2 was from Statens Serum Institut (Ref 3459).

### Biofilm inhibition tests

Overnight cultures were adjusted to an OD_600nm_ of 0.05 before inoculating 50 µl into 96-well polyvinyl chloride (PVC) plates (Falcon; Becton Dickinson Labware, Oxnard, CA) and added at a 1:1 ratio to 50 µl of filter-sterilized and purified polysaccharide at the desired concentration: i) 100 µg/ml for standardized antibiofilm activity experiments; ii) 3.125-100 µg/ml for test level of activity of the macromolecules. Biofilms were left to grow for 16 h at 37°C before quantification as described in (7).

### Polysaccharide purification

G2cps polysaccharide was obtained from 24h cultures as described in (7). Briefly, CFT073 *E. coli* strains were grown in M63B1 0.4%glucose for 24h at 37°C. After cold centrifugation at 7500 rpm for 10 minutes, supernatants were filtered through a 0.22µm filter. Polysaccharides were precipitated from cell-free supernatant using ice-cold ethanol (3:1 ratio ethanol / supernatant) followed by centrifugation for 2h at 10000 rpm and 4°C. Precipitated polysaccharides were resuspended and dialyzed against deionized water (10 kDa cassettes; Pierce, Rockford, IL). Total polysaccharide concentrations were measured by Duvois colorimetric assay (42). The polysaccharides were finally separated from residual lipopolysaccharides by gel filtration on Sepharose 6BCL column (GE Healthcare Life Sciences) in 1% sodium deoxycholate (43) and dualized against deionized water. Polysaccharide concentration was determined using High-Performance Anion-Exchange Chromatography with Pulsed Amperometry Detection (HPAEC-PAD) (Thermo Fischer Scientific, Dionex, Sunnyval, CA) as previously described(44).

### Polysaccharide structure analysis by High Performance Anion Exchange Chromatography with Pulsed Amperometric Detection (HPAEC-PAD)

Polysaccharide G2cps was hydrolyzed with hydrofluoric acid (HF) (48% by mass) for 16h at room temperature. HF was removed by drying under a stream of nitrogen at 40°C. The sample was redissolved in water and then hydrolyzed with 2M trifluoroacetic acid (TFA) for 2h at 121°C. TFA was removed by drying under a stream of nitrogen at 40°C. The sample was redissolved in water prior to analysis. For the sake of comparison, both HF-alone and TFA-alone hydrolysis were also performed on polysaccharide G2cps. Pulsed amperometric detection was used incorporating a quadruple-potential waveform. Data were collected and analyzed on computers equipped with Dionex Chromeleon software (Dionex). Commercial monosaccharides were used as standards.

### Nuclear Magnetic Resonance (NMR) spectroscopy

Approximately 5 mg of polysaccharide was lyophilized once, dissolved in 700 μl of deuterated water and 550 μl were introduced in a 5mm tube. NMR spectra were collected on a Bruker Avance 500MHz spectrometer running Topspin 2.1 software at an indicated probe temperature of 20°C. Spectra were measured for solutions in D2O with 0.01% DSS as an internal standard for proton (^1^H) NMR (δ 0.00 ppm). Phosphoric acid (2%) was used as an external standard for phosphorus (^31^P) NMR (δ 0.00 ppm) and TSP as an external standard for carbone (^13^C) NMR (δ 0.00 ppm). The pulse programs used were those in the Bruker library except for the correlation spectroscopy (COSY) and total correlation spectroscopy (TOCSY) diffusion experiments. The diffusion delay was 80 ms for these two-dimensional experiments and the mixing time for the TOCSY was 100 ms.

The two dimensional ^1^H-^31^P experiment was acquired using a *J*_H-P_ coupling value of 7Hz. The ^1^H-^13^C heteronuclear single quantum coherence (HSQC) spectra were acquired using a ^n^*J*_H-C_ coupling value of 145Hz and a long range ^n^*J*_H-C_ coupling value of 5Hz for heteronuclear multiple bond correlation (HMBC).

### High Performance Size Exclusion Chromatography methods with triple detection: *Mw* and [*η*] determinations

Polysaccharides were analyzed at a concentration between 600 to 1000 μg/ml in water. Analyses were performed using a TDA301 Viscotek HPSEC system consisting of an automated sampler with a HPLC pump with an injection loop of 100 μl, an on-line degasser, a viscometer, a refractive index (RI), and a right-angle laser light scattering detector (RALLS). The HPSEC analyses of polysaccharide were performed using TSK G4000 PWXL (0.7×30 cm) and TSK G6000 PWXL analytical columns (0.7×30 cm) connected in series at a flow rate of 0.5 ml/min, at 30°C. Tris buffer 2.5mM, pH 7.5 or phosphate buffer 200mM pH 6.9 was used as mobile phase. The columns were kept at a constant temperature of 30°C. Omnisec software (Malvern) was used for data collection and data processing. This triple detection consisting of online RI, RALLS and viscometric detector was used to determine the molecular weight (*Mw*) and the intrinsic viscosity ([*η*]) of polysaccharides.

### Size reduction of G2cps polysaccharide

75 µg of polysaccharide were hydrolyzed in solution with ascorbic acid, copper and iron sulfate for 2 h at 30°C. Hydrolysis kinetics was followed by HPSEC analysis for 15 min, 1 h and 24 h.

### Contact angle measurements

Glass or polyester plastic microscopy coverslip slides were treated with 80 µl of distilled deionized water or with 100 µg/ml solution of purified active and inactive polysaccharides. Slides were dried under a laminar flow hood and 2.5 μl drops of water were deposited using a Kruss DSA4 Drop Shape Analyzer on the surface of coated and uncoated slides, and contact angle was then measured 3 times.

### Atomic Force Microscopy measurements

Microscopy glass slides were treated with 100 µg/ml solution of purified active and inactive polysaccharides for 10 minutes and rinsed with ultrapure water. AFM measurements were performed in ultrapure water, using a Fastscan Dimension Icon with Nanoscope V controller (Bruker). Images were obtained in Peak Force tapping mode, using gold-coated silicon nitride tips (NPG, Bruker), with a maximum applied force of 500 pN, a scan rate of 1 Hz, a peak force amplitude of 300 nm and a peak fore frequency of 2 kHz. For quantifying coated-surface hydrophobicity by means of chemical force microscopy, hydrophobic tips were prepared by immersing gold-coated silicon nitride tips (NPG, Bruker) for 12h in 1 mM solution of dodecanethiol (Sigma) in ethanol, rinsed with ethanol and dried with N_2_. Spatial mappings were obtained by recording 32 × 32 approach-retract force-distance curves on 5 µm × 5 µm areas, with a maximum applied force of 500 pN and an approach and retraction speed of 500 nm/s.

### Electrokinetic measurements

1 g/L stock suspensions of macromolecules were first prepared in ultrapure water from corresponding frozen powders and, after 48 h, suspensions were diluted at 250 mg/L and adjusted to pH 5. For each macromolecule tested, batches were then prepared in order to obtain a series of suspensions with NaNO_3_ background electrolyte concentration in the range 1 to 100 mM. Final macromolecule concentration and pH conditions were adopted to optimize the measured electrophoretic response after testing a range of particle concentrations from 50 to 250 mg/L and four pH conditions (3, 5, 7 and 9). All dispersions, stock and diluted suspensions were stored at 4°C and were re-acclimatized at room temperature prior to measurements.

24 to 48h after batches preparation, diffusion coefficients and electrophoretic mobilities of the bacterial polysaccharides were measured by Dynamic Light Scattering (DLS, Supporting Table S2) and Phase Analysis Light Scattering (Malvern Instruments). Each reported data point in Figure 4 corresponds to three measurements performed on 3 different aliquots of a given batch macromolecule dispersion.

### Quantitative assessment of polysaccharide electrophoretic mobility

Electrophoretic mobility of the various macromolecules of interest were measured as a function of NaNO_3_ electrolyte concentration (Figure 4) and were collated with the analytical theory developed by Ohshima for electrophoresis of soft colloids (33) in the Hermans-Fujita limit applicable to soft polyelectrolytes (34). According to the latter, μ is defined by the expression

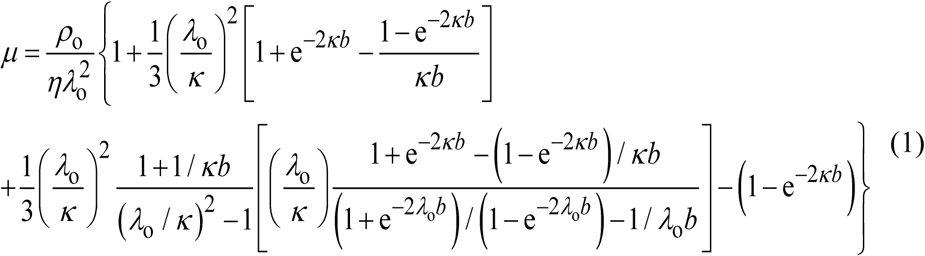

, where *ρ*_*o*_ represents the density of negative charges carried by the macromolecule, *κ* is the reciprocal Debye layer thickness defined by *κ* = [2*F* ^2^*c*_0_/ (*εRT*)] ^1/2^ with *R* the gas constant, *T* the absolute temperature, *F* the Faraday number and *c*_o_ the bulk concentration of a 1:1 electrolyte (NaNO_3_ in this work), *λ*_o_ is the so-called softness parameter with 1/*λ*_o_ corresponding to the characteristic penetration length of the electroosmotic flow within the macromolecule. The quantity *b* in eq 1 is the radius of the polyelectrolyte macromolecule. The set of (*ρ*_o_, 1/*λ*_o_) couples obtained for all polysaccharidic macromolecules are reported in Supporting Table S2 and the required values of *b* for data fitting to eq 1 were determined by Dynamic Light Scattering and are also given in Supporting Table S2.

### Statistical analysis

Two-tailed unpaired *t-test* with Welch correction analyses were performed using Prism 9.0 for Mac OS X (GraphPad Software). Each experiment was performed at least three times. All data are expressed as mean (± standard deviation, SD) in figures. Differences were considered statistically significant for *P* values of <0.05; * p<0.05; ** p<0.01; *** p<0.001. **** p<0.0001.

## ACKNOWLEDGEMENTS

We thank Olaya Rendueles for helpful discussions and Nadia Izadi and Laurence Mulard for critical reading of the manuscript. We are grateful to Heike Claus, Ulrich Vogel and Muhamed-kheir Taha, for generously providing us with assistance with some of the strains used in this study. This work was supported by a collaborative research grant Institut Pasteur and Sanofi, by grants from the French Government’s Investissement d’Avenir program, Laboratoire d’Excellence “Integrative Biology of Emerging Infectious Diseases” (grant n°ANR-10-LABX-62-IBEID) and by the Fondation pour la Recherche Médicale (grant DEQ20180339185). J. B-B was the recipient of a long-term post-doctoral fellowship from the Federation of European Biochemical Societies (FEBS) and by the European Union’s Horizon 2020 research and innovation program under the Marie Skłodowska-Curie grant agreement No 842629. This work was partly carried out in the Pôle de compétences Physico-Chimie de l’Environnement, LIEC laboratory UMR 7360 CNRS -Université de Lorraine.

