## Supplementary Tables S1 to S3 and Figures S1 to S8 for "Bacterial capsular polysaccharides with antibiofilm activity share common biophysical and electrokinetic properties"

Joaquín Bayard-Bernal et *al.*

**This PDF file includes:**

Supporting Tables S1 to S3

Supporting Figures S1 to S8

### SUPPORTING TABLES

**Supporting Table S1.** Name, origin, primary structure and properties of the polysaccharides used in this study[1]

| Name[1] | Bacteria | Primary structure | Antibiofilm activity | References |
|---|---|---|---|---|
| PnPS 1 | *Streptococcus pneumoniae* serotype 1 | 2/3-(OAc)$_{68\%}$<br>\|<br>[3)-α-AAT*p*-(1→4)-α-D-Gal*p*A-(1→3)-α-D-Gal*p*A-(1→]$_n$ | - | (1) (2) |
| PnPS 2 | *Streptococcus pneumoniae* serotype 2 | [4)-β-D-Glc*p*-(1→3)-α-L-Rha*p*-(1→3)-α-L-Rha*p*-(1→3)-β-L-Rha*p*-(1→]$_n$<br>2<br>↑<br>1<br>α-D-GlcA*p* -(1→6)- α-D-Glc*p* | - | (3) |
| PnPS 3 | *Streptococcus pneumoniae* serotype 3 | [4)-β-D-Glc*p*-(1→3)-β-D-Glc*p*A-(1→]$_n$ | **G+ / ~ G-** | (4) (2) |
| PnPS 4 | *Streptococcus pneumoniae* serotype 4 | [3)-β-D-Man*p*NAc-(1→3)-α-L-Fuc*p*NAc-(1→3)-α-D-Gal*p*NAc-(1→4)-α-D-Gal*p*-(2,3-(s)-pyruvate)(1→]$_n$ | - | (5) (6) |
| PnPS 5 | *Streptococcus pneumoniae* serotype 5 | [4)-β-D-Glc*p*-(1→4)-α-L-Fuc*p*NAc-(1→3)-β-D-Sug*p*-(1→]$_n$<br>3<br>↑<br>1<br>α-L-Pne*p*NAc-(1→2)-β-D-Glc*p*A | - | (7) (6) |
| PnPS 6B | *Streptococcus pneumoniae* serotype 6B | [2)-α-D-Gal*p*-(1→3)-α-D-Glc*p*-(1→3)-α-L-Rha*p*-(1→4)-D-Ribitol-5-*P*-(O→]$_n$ | - | (8) (2) |
| PnPS 7F | *Streptococcus pneumoniae* serotype 7F | 2-OAc<br>\|<br>[6)-α-D-Gal*p*-(1→3)-β-L-Rha*p*-(1→4)-β-D-Glc*p*-(1→3)-β-D-Gal*p*NAc-(1→]$_n$<br>2 ↑ 1 β-D-Gal*p* (at Gal*p*); 4 ↑ 1 α-D-Glc*p*NAc-(1→ 2)-α-L-Rha*p* (at Gal*p*NAc) | - | (9) (2) |
| PnPS 8 | *Streptococcus pneumoniae* serotype 8 | [4)-β-D-Glc*p*A-(1→4)-β-D-Glc*p*-(1→4)-α-D-Glc*p*-(1→4)-α-D-Gal*p*-(1→]$_n$ | - | (10) (2) |
| PnPS 9N | *Streptococcus pneumoniae* serotype 9N | [4)-α-D-Glc*p*A-(1→3)-α-D-Glc*p*-(1→3)-β-D-Man*p*NAc-(1→4)- β-D-Glc*p*-(1→4)-α-D-Glc*pNAc*-(1→]$_n$ | - | (11) (2) |
| PnPS 9V | *Streptococcus pneumoniae* serotype 9V | [4)-α-D-Glc*p*-(1→4)-α-D-Glc*p*A-(1→3)-α-L-Gal*p*-(1→3)-β-D-Man*p*NAc-(1→4)-β-D-Glc*p*-(1→]$_n$<br>\| 2/3-(OAc)$_{80\%}$ (at Glc*p*A); \| 4/6-(OAc)$_{113\%}$ (at Man*p*NAc) | - | (12) (2) |

| | | | | |
|---|---|---|---|---|
| PnPS 10A | *Streptococcus pneumoniae* serotype10A | β-D-Gal*p*<br>1<br>↓<br>6<br>[5)-β-D-Gal*f*-(1→3)-β-D-Gal*p*-(1→4)-β-D-Gal*pNAc*-(1→3)-α-D-Gal*p*-(1→2)-D-Ribitol-5-*P*-(O→]$_n$<br>3<br>↑<br>1<br>β-D-Gal*f* | - | (13) (2) |
| PnPS 11A | *Streptococcus pneumoniae* serotype 11A | 2/3-(OAc)$_{110\%}$ 4/6-(OAc)$_{50\%}$<br>I I<br>[6)-α-D-Glc*p*-(1→4)-α-D-Gal*p*-(1→3)-β-D-Gal*p*-(1→4)-β-D-Glc*p*-(1→]$_n$<br>4<br>I<br>*O*)-*P*-1-Glycerol | - | (14) (2) |
| PnPS 12F | *Streptococcus pneumoniae* serotype 12F | [4)-α-L-Fuc*p*NAc-(1→3)-β-D-Gal*p*NAc-(1→4)-β-D-Man*p*NAcA-(1→]$_n$<br>3 3<br>↑ ↑<br>1 1<br>α-D-Gal*p* α-D-Glc*p*-(1→2)-α-D-Glc*p* | **G- only** | (15) (2) |
| PnPS 14 | *Streptococcus pneumoniae* serotype 14 | [6)-β-D-Glc*pN*Ac-(1→3)-β-D-Gal*p*-(1→4)-β-D-Glc*p*-(1→]$_n$<br>4<br>↑<br>1<br>β-D-Gal*p* | - | (16) (2) |
| PnPS 15B | *Streptococcus pneumoniae* serotype15B | [6)-β-D-Glc*pN*Ac-(1→3)-β-D-Gal*p*-(1→4)-β-D-Glc*p*-(1→]$_n$<br>4<br>↑<br>1<br>α-D-Gal*p*-(1→ 2)- β-D-Gal*p*<br>I 3<br>2/3/4/6-(OAc)$_{85\%}$ I<br>*O)*-*P*-2-Glycerol | - | (17) (6) |
| PnPS 17F | *Streptococcus pneumoniae* serotype 17F | 2-OAc<br>I<br>[3)-β-L-Rha*p*-(1→4)-D-β-D-Glc*p*-(1→3)-α-D-Gal*p*-(1→3)-β-L-Rha*p*-(1→4)-α-L-Rha*p*-(1→2)-D-Ara-ol-1-*P*-(O→]$_n$<br>4<br>↑<br>1<br>β-D-Gal*p* | - | (18) (2) |
| PnPS 18C | *Streptococcus pneumoniae* serotype 18C | 6-(OAc)$_{30\%}$<br>I<br>α-D-Glc*p*<br>1<br>↓<br>2<br>[4)-β-D-Glc*p*-(1→4)-β-D-Gal*p*-(1→4)-α-D-Glc*p*-(1→3)-α-L-Rha*p*(1→]$_n$<br>3<br>I<br>*O)*-*P*-2-Glycerol | **G+ only** | (19) (6) |
| PnPS 19A | *Streptococcus pneumoniae* serotype 19A | [4)-β-D-Man*p*NAc-(1→4)-α-D-Glc*p*-(1→3)-α-L-Rha*p*-1-*P*-(O→]$_n$ | - | (20) (2) |
| PnPS 19F | *Streptococcus pneumoniae* serotype 19F | [4)-β-D-Man*p*NAc-(1→4)-α-D-Glc*p*-(1→2)-α-L-Rha*p*-1-*P*-(O→]$_n$ | - | (21) (2) |
| PnPS 20 | *Streptococcus pneumoniae* serotype 20 | 6-(OAc)$_{90\%}$<br>I<br>[6)-α-D-Glc*p*-(1→6)-β-D-Glc*p*-(1→3)-β-D-Gal*f*-(1→3)-β-D-Glc*p* -(1→3)-α-D-Glc*pN*Ac-1-*P*-(O→]$_n$<br>I 4<br>5-(OAc)$_{90\%}$ ↑<br>1<br>β-D-Gal*f*<br>I<br>2-(OAc)$_{90\%}$ | - | (12) (2) |
| PnPS 22F | *Streptococcus pneumoniae* serotype 22F | β-D-Glc*p*<br>1<br>↓<br>3<br>[4)-β-D-Glc*p*A-(1→4)-β-L-Rha*p*-(1→4)-α-D-Glc*p*-(1→3)-α-D-Gal*f*-(1→2)-α-L-Rha*p*-(1→]$_n$<br>I<br>2-(OAc)$_{80\%}$ | - | (22) (2) |
| PnPS 23F | *Streptococcus pneumoniae* serotype 23F | α-L-Rha*p*<br>1<br>↓<br>2<br>[4)-β-D-Glc*p*-(1→4)-β-D-Gal*p*-(1→4)-β-L-Rha*p*-(1→]$_n$<br>3<br>I<br>*O)*-*P*-2-Glycerol | - | (23) (2) |
| PnPS 33F | *Streptococcus pneumoniae* serotype 33F | [3)-β-D-Gal*p*-(1→3)-α-D-Gal*p*-(1→3)-β-D-Gal*f*-(1→3)-β-D-Glc*p* -(1→5)-β-D-Gal*f*(1→]$_n$<br>2 I<br>↑ 2-(OAc)$_{50\%}$<br>1<br>α-D-Gal*p* | - | (24) (2) |
| CWPS | noncapsulated *Streptococcus pneumoniae* strain CSR SCS2 | [6)-β-D-Glc*p*-(1→3)-α-AAT*p*-(1→4)-α-D-Gal*p*NAc-(1→3)-β-D-Gal*p*NAc-(1→1)-D-Ribitol-5-*P*-(O→]$_n$<br>6<br>I<br>O)-*P*-Cho | - | (25) |

| | | | | |
|---|---|---|---|---|
| Vi | *Salmonella enterica Serovar Typhi* | 3-(OAc)$_{90\%}$<br>I<br>[4)-α-D-Gal*p*NAcA-(1→]$_n$ | G+ / ~ G- | (26) |
| PRP | *Haemophilus influenzae* serotype b | [3)-β-D-Rib*f*-(1→1)-D-Ribitol-5-*P*-(O→]$_n$ | G+ / ~ G- | (27) |
| MenA | *Neisseria meningitidis* serogroup A | 3/4-(OAc)$_{90\%}$<br>I<br>[6)-α-D-Man*p*NAc-1-*P*-(O→]$_n$ | G+ & G- | (28) |
| MenC | *Neisseria meningitidis* serogroup C | 7/8-(OAc)$_{90\%}$<br>I<br>[9)-α-D-Neu*p5Ac*-(2→]$_n$ | G+ & G- | (28) |
| MenY | *Neisseria meningitidis* serogroup Y | 7/9-(OAc)$_{\sim 50\%}$<br>I<br>[4)-α-D-Neu*p*5Ac-(2→6)-α-D-Glc*p*-(1→]$_n$ | G+ & G- | (28) |
| MenW135 | *Neisseria meningitidis* serogroup W135 | 7/9-(OAc)$_{\sim 60\%}$<br>I<br>[4)-α-D-Neu*p*5Ac-(2→6)-α-D-Gal*p*-(1→]$_n$ | G+ & G- | (28) |
| G2cps | *Escherichia coli* strain CFT073 | 3-(OAc)$_{100\%}$<br>I<br>[4)-α-D-Gal*p*-(1→2)-Glycerol-3-*P*-(O→]$_n$ | G+ & G- | (29), this study |

*In red:* active antibiofilm polysaccharides.

[1]: All polysaccharides produced and purified from *S. pneumoniae*, *N. meningiditis*, *S.* Typhi and *H. influenzae* strains used in this study were obtained from Sanofi, Marcy l'Etoile France and Swiftwater, USA.

**Supporting Table S2**. Electrokinetic and size properties of the polysaccharidic macromolecules tested in this work.

| **Polysaccharide** | $\lvert \rho_0/F \rvert$ **(mm)** | **$1/\lambda_0$ (nm)** | **Size *d* (nm) ± standard deviation** |
|---|---|---|---|
| Vi | | | 15.6 ± 0.7 |
| PnPs19A | 15.7 | 2.8 | 20.9 ± 4.2 |
| PnPS8 | 20.5 | 2.5 | 23.4 ±1.4 |
| MenA | | | 12.1 ± 0.7 |
| PnPS3 | 18.7 | 3.2 | 26.3 ± 3.9 |
| PnPS1 | 20.1 | 2.4 | 26 ± 2.3 |
| PnPS9V | 25.7 | 1.4 | 8.7 ± 1.6 |
| PRP | 20.3 | 3.3 | 28.4 ± 5 |
| G2cps | 18.1 | 3.2 | 40±15 |
| PnPS7F | 10.8 | 3.1 | 39.4 ± 2.7 |
| PnPS22F | 15.5 | 2.3 | 26.3 ± 1.9 |
| PnPS14 | 30.3 | 1.6 | 15.2 ±5.1 |
| MenW135 | | | 28.3 ± 3.1 |
| PnPS18C | | | 26.7 ± 2.2 |
| PnPS12F | | | 29.5 ± 3.3 |

In red: active broad-spectrum antibiofilm polysaccharides. In blue: active narrow spectrum antibiofilm polysaccharides. *d* stands for the diameter of the macromolecules (*d*=2*b* where *b* is the particle radius involved in eq. 1). The values $\rho_0/F$ and $1/\lambda_0$ for the polysaccharides Vi, MenA, MenW135 and PnPS18C, PnPS12F fall within the space solution defined by the red and blue zones in Figure 5, respectively.

#### Supporting Table S3. Bacterial strains used in this study.

| Strain | Relevant characteristics | Source/Reference |
|---|---|---|
| *Strains producing antibiofilm polysaccharides* | | |
| *E. coli* CFT073 | Uropathogenic *E.coli* - group 2 capsule producer | (1) |
| *Strains used as biofilm test panel* | | |
| *E. coli* K-12 MG1655 *F'tet ΔtraD* | *traD*::apra plasmid, $Apra^R$, $Tet^R$ | (2) |
| *Enterobacter cloacae 1092* | | (3) |
| *Klebsiella pneumoniae* U21 | Clinical isolate | (3) |
| *Staphylococcus aureus* 15981 | | (4) |
| *Staphylococcus epidermidis* 047 | | (5) |

### SUPPORTING FIGURES

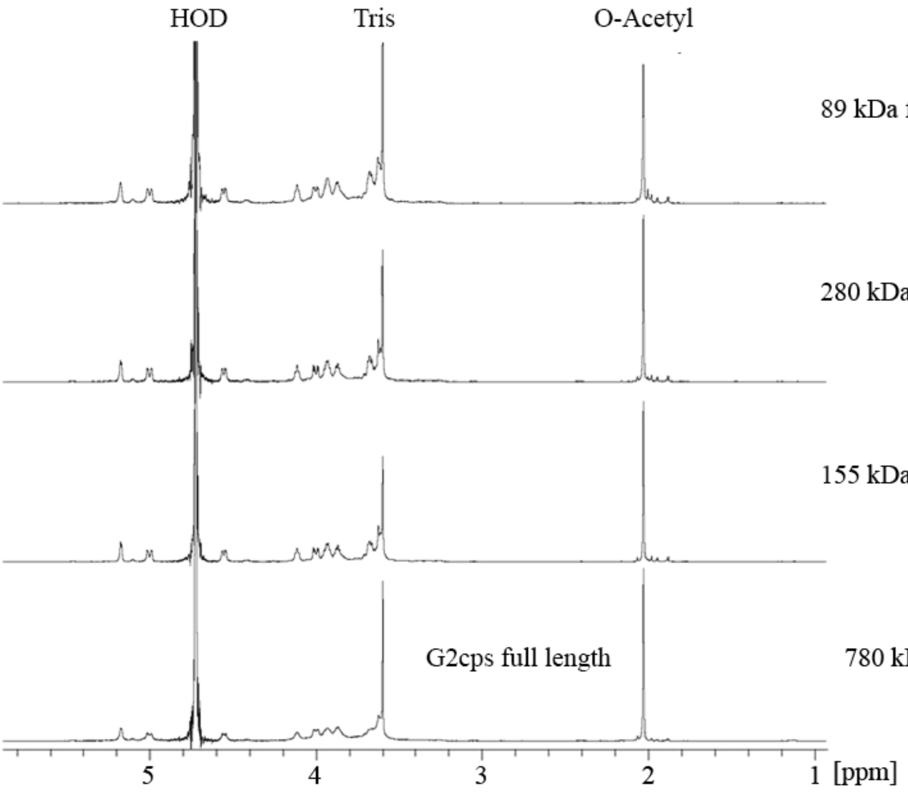


Supporting Figure S1**. $^1$H NMR spectra of native G2cps polysaccharide and corresponding fragments obtained by radical oxidation hydrolysis**. The analysis was performed at 20°C). On each spectrum is indicated the *Mw* (kDa) determined by HPSEC. HOD: signal of residual water.

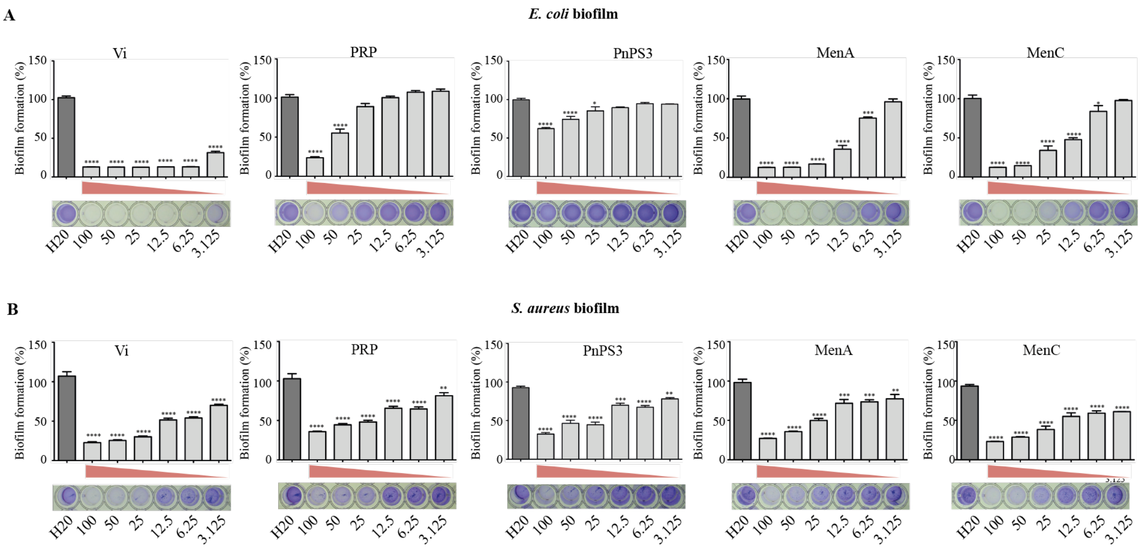


Supporting Figure S2. **Level of antibiofilm activity of bacterial polysaccharides.** *E. coli* **(A)** and *S. aureus* **(B)** biofilm inhibition test (CV staining in microtiter plates) in presence of increasing concentrations of the indicated bacterial capsular polysaccharides produced by *Streptococcus pneumoniae*, *Salmonella enterica* serovar Typhi*, Haemophilus influenzae, Neisseria meningitidis.* Polysaccharide concentrations range from 3.125 to 100 µg/ml. Distilled water was used as a negative control. Each experiment was performed at least 3 times. * $p<0.05$; ** $p<0.01$; *** $p<0.001$; **** $p<0.0001$.

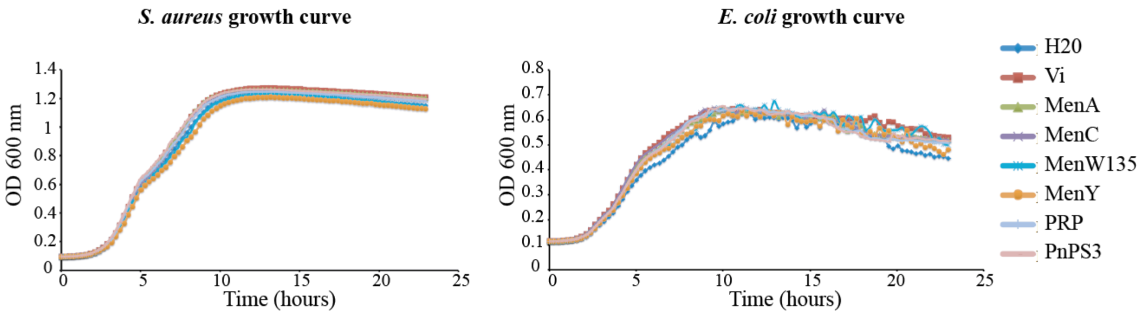


Supporting Figure S3. **The identified antibiofilm polysaccharides are non-biocidal.** Growth curves of *S. aureus* and *E. coli* exposed to 100 µg/ml of each indicated tested macromolecule. The bacterial strains were inoculated in microtiter plates at $OD_{600nm}$ of 0.05 and let to grow with agitation for 24 h at 37ºC. Bacterial growth was determined using a TECAN plate reader.

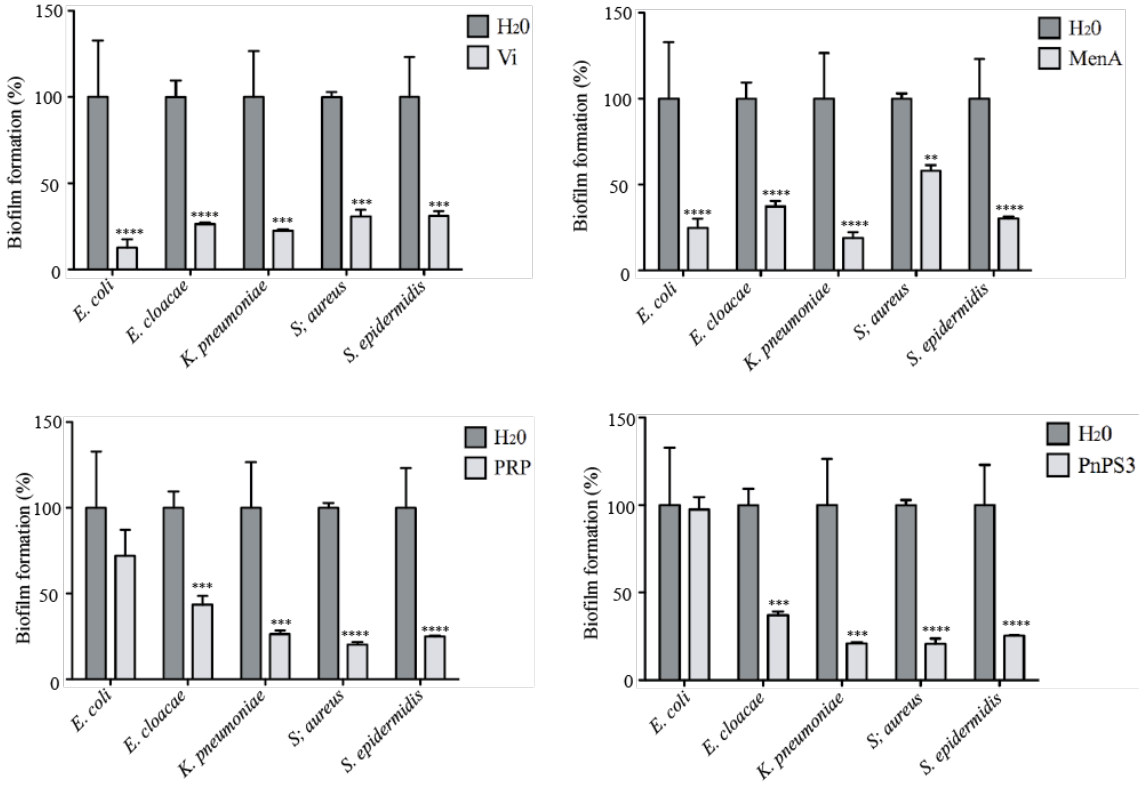


Supporting Figure S4. **Spectrum of activity of several active polysaccharides.** Antibiofilm activity of purified polysaccharides Vi, MenA, PRP and PnPS3 was assessed over a panel of relevant biofilm-forming pathogenic bacteria. Tested Gram-positive bacteria include *Staphylococcus aureus* 15981 and *Staphylococcus epidermidis* 0-47. Tested Gram-negative bacteria include *Escherichia coli*, *Enterobacter cloacae* 1092 and *Klebsiella pneumoniae* 21. Biofilm inhibition tests were performed in presence of 50 µg/ml of polysaccharide. Distilled water was used as a negative control. Each experiment was performed in triplicate. ** p<0.01; *** p<0.001. **** p<0.0001.

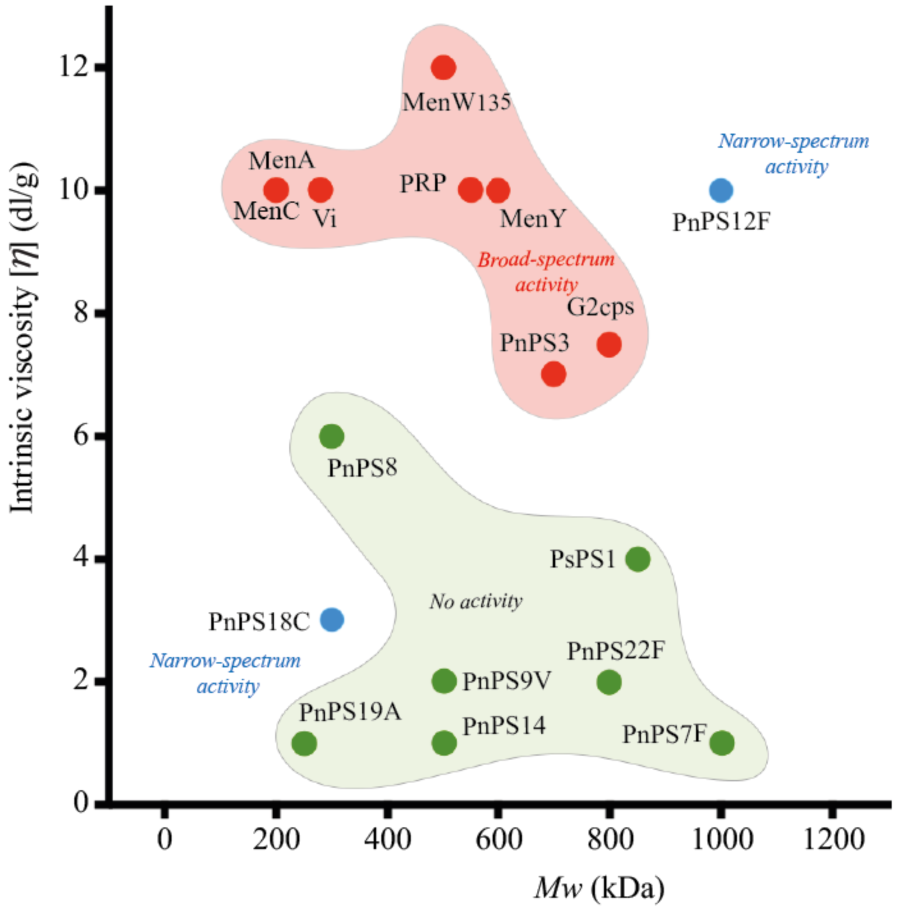


Supporting Figure S5. **Classification of the tested polysaccharides according to their molecular weight (*Mw*) and intrinsic viscosity [*η*].** In red: broad spectrum active macromolecules. In green: inactive macromolecules. In blue: polysaccharides with narrow spectrum antibiofilm activity

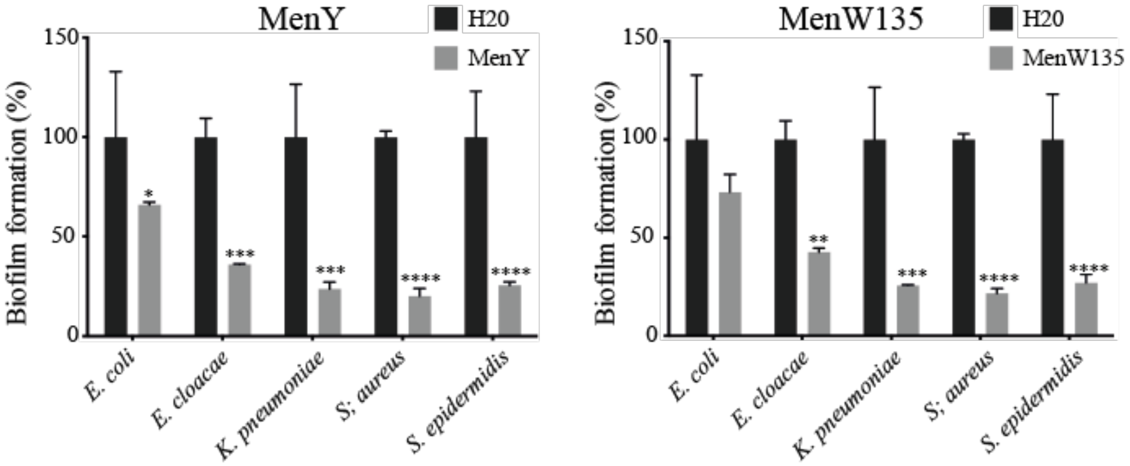


Supporting Figure S6. **Spectrum of action of MenY and MenW135 over a panel of relevant biofilm-forming pathogenic bacteria**. Tested Gram+ bacteria include *Staphylococcus aureus* 15981 and *Staphylococcus epidermidis* 047. Tested Gram- bacteria include *Escherichia coli* K12 MG1655 carrying the F plasmid, *Enterobacter cloacae* 1092 and *Klebsiella pneumoniae* 21. Biofilm inhibition tests were performed in presence of 50 μg/ml of polysaccharide. Distilled water was used as a negative control. Each experiment was performed in triplicate. * $p<0.05$; ** $p<0.01$; *** $p<0.001$; **** $p<0.0001$.

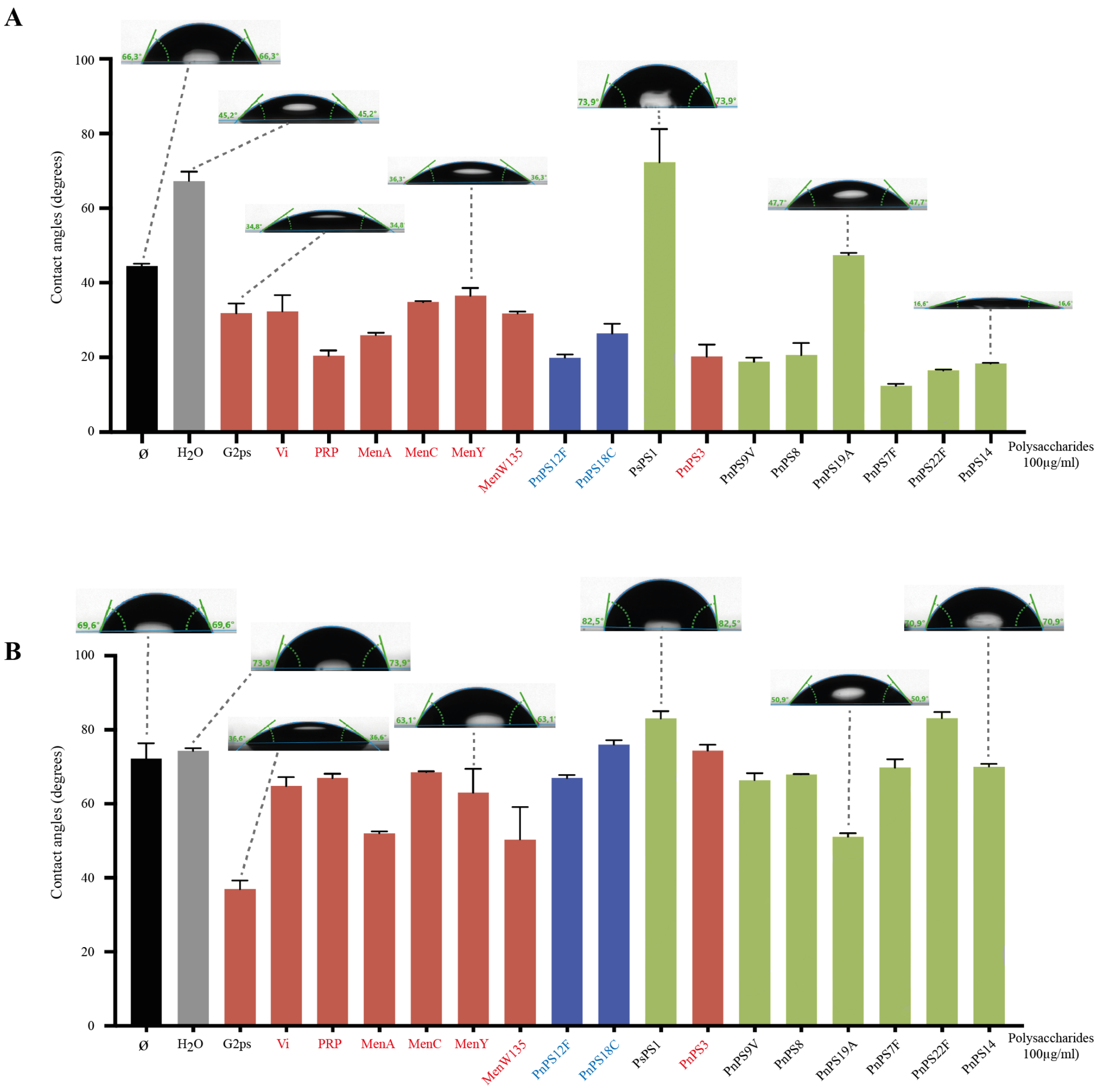


Supporting Figure S7. **Determination of surface contact angle of a drop of water on glass or plastic surfaces treated with active and inactive polysaccharides**. Glass (**A**) and polyester plastic (**B)** microscopy slides untreated, treated with $H_2O$ or treated with active broad-spectrum activity polysaccharides (in red), active narrow-spectrum activity polysaccharides (in blue) and inactive polysaccharides (in green) (see Table 1) in presence of 100 µg/ml of polysaccharide. Representative pictures of water droplets used to determine contact angle are presented.

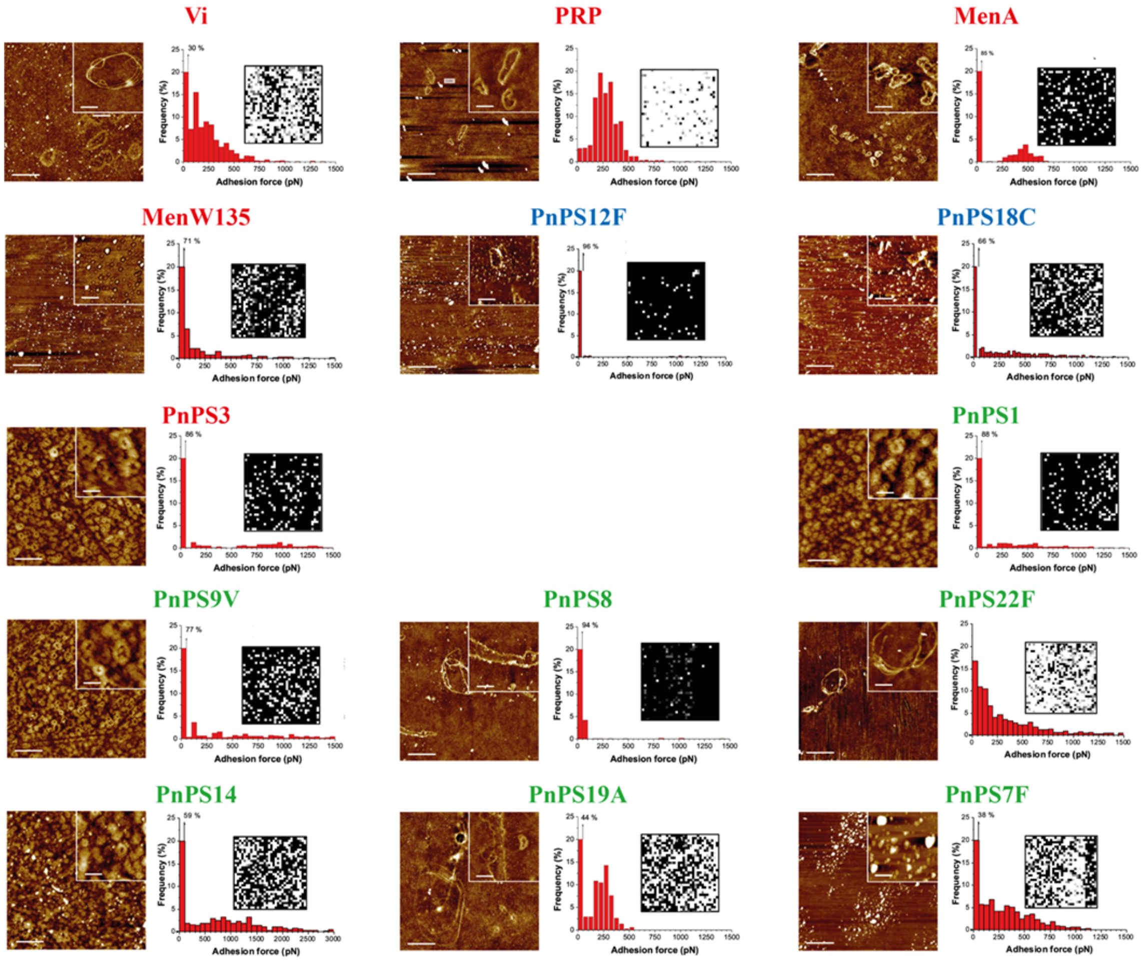


Supporting Figure S8. **Representative AFM peak force images and adhesion force histograms of selected active and inactive polysaccharides.** Each couple of panels correspond to a given polysaccharide (specified). In each couple of panels, AFM images are on the left and adhesion force histograms on the right. Polysaccharides with broad-spectrum activity (indicated in red), narrow-spectrum activity polysaccharides (indicated in blue) and inactive polysaccharides (indicated in green) were deposited on a glass surface at 100 µg/mL and imaged in ultrapure water. Scale bars correspond to 1 µm. In each image, the inset shows the sample at higher magnification (scale bars = 250 nm). Adhesion force histograms (n=1024 force-distance curves) and corresponding adhesion force maps were recorded between AFM hydrophobic $CH_3$-tips and glass surfaces coated with selected active and inactive polysaccharides. Each map has been recorded on a 5 µm x 5 µm area. Grey scale: 150 pN, black pixels correspond to non-adhesive events.
